# Evaluation of the effects of SARS-CoV-2 genetic mutations on diagnostic RT-PCR assays

**DOI:** 10.1101/2021.01.19.426622

**Authors:** Takeru Nakabayashi, Yuki Kawasaki, Koichiro Murashima, Kazuya Omi, Satoshi Yuhara

## Abstract

Several mutant strains of severe acute respiratory syndrome coronavirus 2 (SARS-CoV-2) are emerging. Mismatch(es) in primer/probe binding regions would decrease the detection sensitivity of the PCR test, thereby affecting the results of clinical testing. In this study, we conducted an in silico survey on SARS-CoV-2 sequence variability within the binding regions of primer/probe published by the Japan National Institute of Infectious Diseases (NIID) and Centers for Disease Control and Prevention (CDC). In silico analysis revealed the presence of mutations in the primer/probe binding regions. We performed RT-PCR assays using synthetic RNAs containing the mutations and showed that some mutations significantly decreased the detection sensitivity of the RT-PCR assays.

Our results highlight the importance of genomic monitoring of SARS-CoV-2 and evaluating the effects of mismatches on PCR testing sensitivity.

## Introduction

Coronavirus disease 2019 (COVID-19) pandemic is caused by the SARS-CoV-2 virus (1), and the global number of cases has reached 63 million as of December 2020 (2). COVID-19 infection is diagnosed via the detection of SARS-CoV-2 RNA in nasopharyngeal, nasal, or saliva specimens by performing the RT-PCR method with the protocol established by the National Institute of Infectious Diseases (NIID) and Centers for Disease Control and Prevention (CDC) that has been widely used in Japan.

The primers and probes for RT-PCR are designed to detect the conserved region of the SARS-CoV-2 RNA sequence. Hence, it is crucial to assess the impact of gene mutations observed in primer/probe binding sites on the sensitivity of SARS-CoV-2 detection. Several in silico surveys have shown the emergence of mutant strains that exhibit mismatches in the primer/probe binding regions; however, these studies did not assess the effect of such mutations on PCR testing (3, 4).

Here, we conducted an in silico survey of sequence variability within the binding regions of primers/probes used in the NIID and CDC protocols and evaluated the detection sensitivity of RT-PCR performed using synthetic RNAs containing frequently observed mutations. We showed that certain primer/probe-template mismatches significantly decreased the sensitivity of RT-PCR assays. Our survey suggests the necessity of monitoring mutations in the viral genome sequence under in silico conditions and evaluating the impact of mutations on diagnosis sensitivity to avoid false negatives.

## Materials and Methods

The whole-genome sequence data of SARS-CoV-2 were downloaded from the GISAID database (July 6, 2020) (5). Genome data with the total length comprising less than 29,000 bases and derived from non-human hosts were excluded (59,621 sequences in total). The region spanning from 27,500th to 29,500th base pairs of each sequence containing the amplification region was extracted, and the sequences that contained N in this region were filtered out (47,836 sequences in total). We aligned the primer and probe sequences developed by NIID and CDC (Table 1) against the nucleotide sequences using glsearch36 (version 36.3.8g) (6). The frequency of occurrence of mismatch between primer and probe sequences was calculated. For each amplification region, we selected the three most frequently observed sequences, in addition to the sequences with mutations at the 3’ end of the primer binding sites. Oligo DNA sequences with these mutations or those identical to the reference sequence (NC_045512.2) (1) were synthesized using GeneArt Strings DNA Fragments (Thermo Scientific). For NIID_N1, CDC_N1, and CDC_N2, the oligos with 150 bp upstream and downstream sequences of the amplification regions were synthesized. For NIID_N2, the oligos with 76 bp upstream and 150 bp downstream sequences of the amplification regions were synthesized owing to the palindromic sequences observed at approximately 80 bp upstream of the amplification region affecting the oligo synthesis. *In vitro* transcription was performed with the synthesized oligos using the CUGA in vitro transcription kit (Nippon Gene, Tokyo, Japan), and the synthetic RNA was purified using RNAclean XP (Beckman Coulter, CA, USA). The synthetic RNA was quantified using NanoDrop (Thermo Scientific) and analyzed using Tapestation (Agilent Technologies). A total of 10,000 copies of synthetic RNA were used in the assay. RT-PCR was performed according to the manufacturer’s instructions or the manual provided by NIID (https://www.niid.go.jp/niid/images/epi/corona/2019-nCoVmanual20200217-en.pdf) using the THUNDERBIRD Probe One-step qRT-PCR Kit (Toyobo).

**Table 1.**
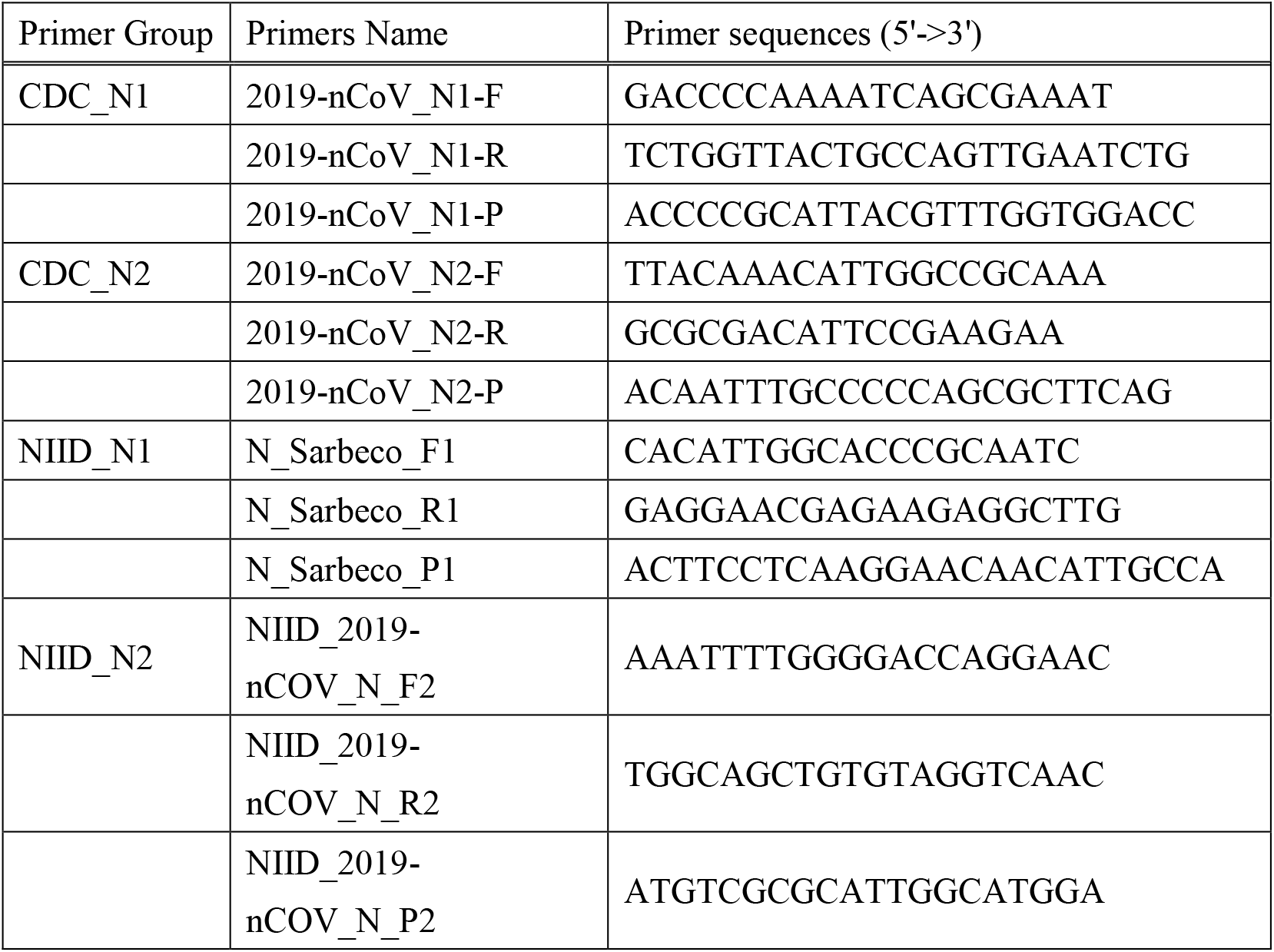
Experimentally evaluated primer and probe sequences analyzed in this study.

**Table 2.** Effects of mismatches in synthetic RNAs on RT-PCR sensitivity.

|  | **Templates** | **Average Ct value** | **ΔWuhan** |
| --- | --- | --- | --- |
| **NIID_N1** | Wuhan-Hu-1 | 29.41 | - |
|  | No.1 | 29.21 | -0.19 |
|  | No.2 | 29.66 | 0.26 |
|  | No.3 | 32.47 | 3.07 |
|  | No.4* | 32.17 | 2.77 |
|  | No.5* | 34.46 | 5.06 |
| **NIID_N2** | Wuhan-Hu-1 | 25.21 | - |
|  | No.1* | Undetermined | >14 |
|  | No.2 | 30.02 | 4.82 |
|  | No.3 | 25.7 | 0.5 |
| **CDC_N1** | Wuhan-Hu-1 | 24.18 | - |
|  | No.1 | 24.12 | -0.08 |
|  | No.2 | 25 | 0.8 |
|  | No.3 | 24.59 | 0.39 |
|  | No.4* | 24.71 | 0.51 |
| **CDC_N2** | Wuhan-Hu-1 | 24.13 | - |
|  | No.1 | 25.49 | 1.39 |
|  | No.2 | 24.84 | 0.74 |
|  | No.3 | 25.29 | 1.19 |
|  | No.4* | 30.39 | 6.29 |

Each Ct value is the mean value of three technical replicates. $\Delta$ Wuhan indicates the difference in Ct values between the mutated and reference sequences. Asterisks indicate the primers that had mismatched nucleotides at the 3' end of the primer binding sites.

## Results

The alignment between the top three most frequently occurring mutations in the SARS-CoV-2 virus genome and primers/probes from NIID and CDC is shown in Figure 1. The forward primer of CDC_N1 showed one nucleotide mismatch with 1.59% (761/47,836) of viral sequences. The incidence rates of the other mismatches were less than 0.5%, which was set as the threshold for sequencing errors in previous studies (4, 7). The forward primer of NIID_N1 (No.4), the reverse primer of NIID_N1 (No.5), the forward primer of NIID_N2 (No.1), the reverse primer of CDC_N1 (No.4), and the forward primer of CDC_N2 (No.4) had nucleotide mismatches at the 3’ end of the primer binding sites with 0.015% (7/47,836), 0.0021% (1/47,836), 0.17% (83/47,836), 0.0021% (1/47,836), and 0.0063% (3/47,836) of viral sequences, respectively.

**Fig. 1.**
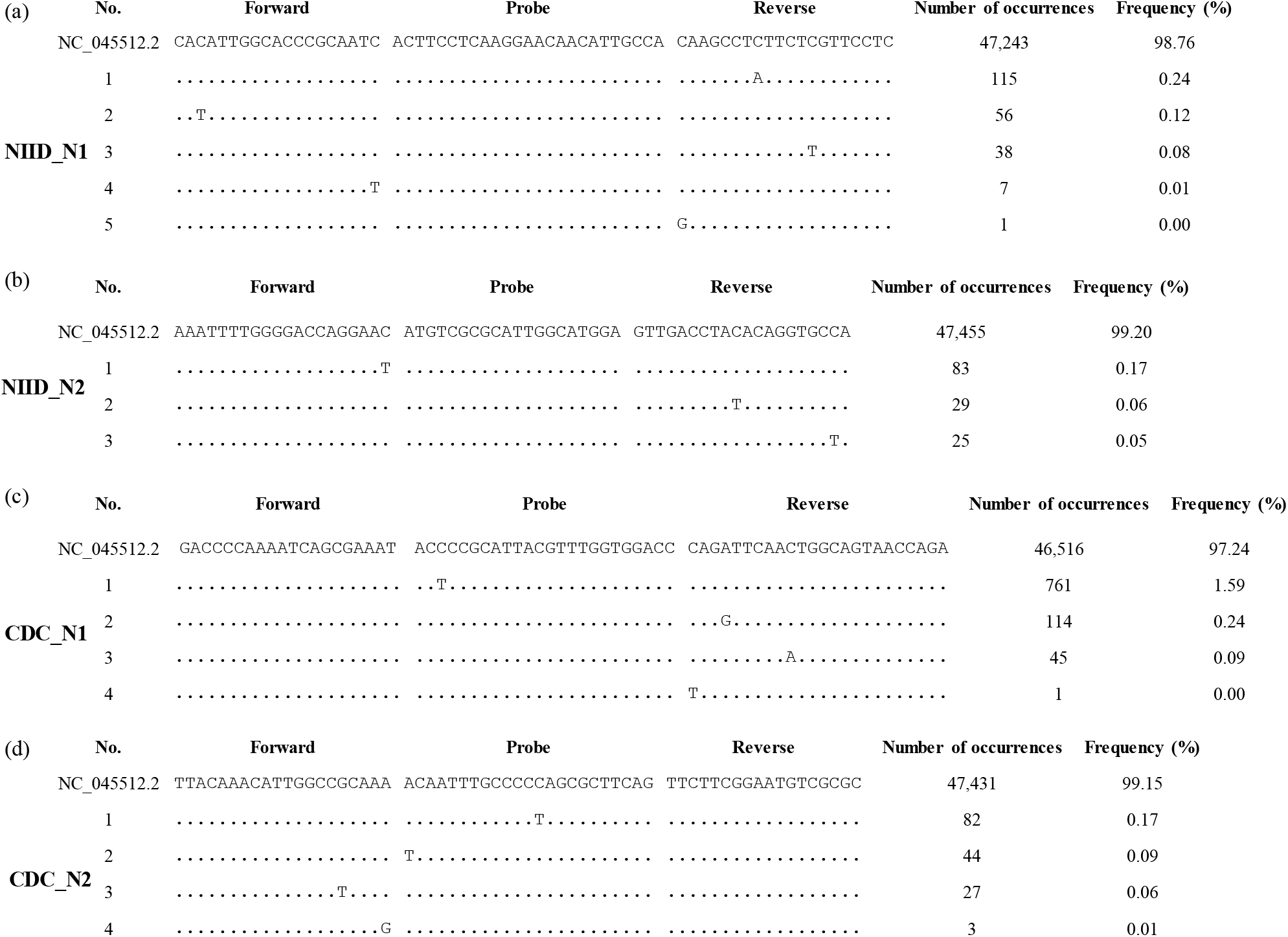
Sequence variants in primers and probe binding regions for NIID_N1 (a), NIID_N2 (b), CDC_N1 (c), and CDC_N2 (d). Sequence variants in 47,836 viral genome sequences aligned to the primer/probe binding regions (5’ to 3’) along with the number of sequence variants and the frequency of each variant in descending order. The dots indicate identical nucleotides with the primers and probes.

Next, we performed RT-PCR assays using synthetic RNA with the mismatches shown in Table 1. As expected, when using the synthetic RNAs with mismatches at the 3’ end of the primer binding site (the forward primer of NIID_N1 (No.4), the reverse primer of NIID_N1 (No.5), and the forward primer of CDC_N2 (No.4)), the Ct value increased (2.77~6.29) compared to that observed when using synthetic RNA with reference sequences. Furthermore, when RNA with a mismatch at the 3’ end of the NIID_N2 primer binding site (No.1) was used, it was not detected by PCR. In contrast, the mismatch in the reverse primer of CDC_N1 (No.4) exerted only minor effects on the Ct value (0.51), even though there was a mismatch at the 3’ end of the primer binding site. For the reverse primer of NIID_N1 (No.3) and the reverse primer of NIID_N2 (No.2), the mismatches in the middle of the primer binding sites had effects on the Ct value (3.07, 4.82, respectively).

## Discussion

In the present study, we conducted an in silico survey of mismatches in the binding regions of primer/probe published by NIID and CDC, which are primarily used in Japan. We also investigated the effects of SARS-CoV-2 genomic mutations on the detection sensitivity of RT-PCR testing. The detection sensitivity of RT-PCR assays decreased with most synthetic RNAs containing mutants with mismatched nucleotides at the 3’ end of the primer binding sites. However, in the case of the reverse primer of CDC_N1, a mismatch at the 3’ end of the primer had little effect on the sensitivity of RT-PCR. Some primer mismatches in the middle of the primer binding regions had certain effects on sensitivity. These results indicated that it is difficult to predict the effects of mismatches on the detection sensitivity of RT-PCR assays using only in silico screening.

In both the CDC and NIID methods, the primer/probe was designed with two different regions of the N gene (NIID_N1 and NIID_N2 for NIID, CDC_N1, and CDC_N2 for CDC) of SARS-CoV-2. At present, no virus strains are known that exhibit mutations in both the NIID_N1 and NIID_N2 regions or both the CDC_N1 and CDC_N2 regions. However, to avoid false-negative diagnoses, it is important to monitor mutations in the viral genome sequence and evaluate the effects of these mutations on the detection sensitivity not only under in silico as well as experimental conditions.

